# Persistence and near persistence via trait evolution: pathways to coexistence

**DOI:** 10.1101/2025.09.23.676445

**Authors:** Swati Patel, Lynn Govaert, Kelsey Lyberger, Victor J. Luque, A. Bradley Duthie, Sébastien Lion

**Affiliations:** Department of Mathematics, Oregon State University Corvallis, OR 97331; Department of Evolutionary and Integrative Ecology, Leibniz Institute of Freshwater Ecology and Inland Fisheries, IGB, Müggelseedamm 301, 12587 Berlin; College of Integrative Sciences and Arts, Arizona State University, Mesa, AZ 85212; Department of Biology Stanford University, Stanford, CA 94305; Department of Philosophy University of Valencia, Av. de Blasco Ibáñez, 30, 46010 València, Spain; Department of Biological and Environmental Sciences, University of Stirling, Stirling, Scotland FK9 4LA; CEFE, CNRS, Univ Montpellier, EPHE, IRD, 1919, route de Mende 34293 Montpellier Cedex 5, France

**Keywords:** modern coexistence theory, coevolution, eco-evolutionary dynamics, persistence theory, character displacement

## Abstract

Modern coexistence theory relies on the invasion criterion: each species must be able to increase when rare in a resident community at ecological equilibrium. However, when considering trait evolution, for example of competing species, this concept must be refined to account for evolution potentially feeding back on population dynamics. We analyze a model of two competing species whose quantitative traits evolve under stabilizing selection, assuming fixed but potentially unequal intraspecific trait variances. Using persistence theory, we derive conditions under which both species persist for all initial trait values, and we introduce the concept of ‘near persistence’, in which coexistence holds for a specific, biologically relevant, subset of initial trait configurations. Our analysis reveals that equal trait variances between species, which is commonly assumed in previous work, yield highly specific outcomes and obscure a broader set of coexistence scenarios that emerge under asymmetric trait variation. We identify fifteen distinct qualitative dynamical regimes as a function of initial conditions and trait variances. Moreover, we show that invasion analysis must often be performed at multiple eco-evolutionary equilibria, as invader dynamics may converge to different trait values depending on the ecological and evolutionary context. We discuss the biological implications of these results and perspectives for future work.

## 1 Introduction

Understanding the origin and maintenance of species diversity is a central goal in ecology and evlutionary biology. In community ecology, a major focus is answering the question of how multiple species persist despite constraints imposed by finite resources and interspecific interactions (Vellend, 2016). Many approaches have been applied to answering this question, including local-stability analysis (e.g., May, 1972; Allesina & Tang, 2012; Grilli et al., 2017; Dougoud et al., 2018), models of spatial heterogeneity (e.g., MacArthur & Wilson, 1967; Hanski & Ranta, 1983; Nee & May, 1992; Chesson, 2000a), or invasibility criteria (e.g., Chesson, 2000b; Barabás et al., 2018; Chesson, 2018).

The invasibility criteria approach underlies modern coexistence theory (henceforth MCT), which has been widely applied to investigating the mechanisms that drive species coexistence (e.g., Chesson, 2000b; Barabás et al., 2018; Chesson, 2018). In this approach, ‘coexistence’ is determined by each species having a positive growth rate when rare, suggesting that all species are expected to persist even if at low densities (Chesson, 2000b). There are multiple mechanisms by which coexistence is facilitated, but these mechanisms can be broadly categorised as stabilising or equalising (Chesson, 2000b; Barabás et al., 2018; Chesson, 2018). Stabilising mechanisms cause species to decrease their own population growth relative to other species (e.g., through intraspecific competition), thereby increasing niche differentiation. Equalising mechanisms decrease differences in species growth rates (e.g., through fluctuating environments). Historically, MCT has focused on pair-wise species interactions (e.g., Chesson & Warner, 1981; Chesson, 1982), or invasion criteria for a focal species against a homogenised stable community (e.g., Li & Chesson, 2016), although the framework can be extended to more complex communities (Ranjan et al., 2024).

Although MCT is a unifying framework for investigating mechanisms of coexistence in community ecology, a critical assumption is that evolutionary dynamics are ignored. However, the nature and magnitude of interspecific interactions depend on potentially evolving species traits. In the past decades, there has been increasing evidence that evolution can occur on timescales rapid enough to affect ecological processes and dynamics, and thus also species interactions (Thompson, 1998; Ellner et al., 2011; Farkas et al., 2013; Brunner et al., 2017; Zamorano et al., 2023). Evolution of species traits can therefore alter the degree to which a focal species affects the density of other species and of itself, which in turn can alter the selective pressure on the trait resulting in further evolution. This interaction between ecological and evolutionary dynamics is also referred to as an eco-evolutionary feedback (Hairston Jr et al., 2005; Hendry, 2016). Consequently, the role of stabilising and equalising mechanisms within a community, and therefore the potential for coexistence, might change due to these eco-evolutionary dynamics. Eco-evolutionary dynamics have been modelled extensively (see e.g., Lion, 2018; Patel et al., 2018; Govaert et al., 2019; Cropp & Norbury, 2021; Barabás et al., 2022b; Weyerer et al., 2023), but integration with MCT remains limited.

Although MCT classically ignores evolution, with the implicit assumption that evolution occurs on a much slower time scale than ecological dynamics, recent attempts to incorporate trait evolution have begun. For example, Pastore et al. (2021) applied an eco-evolutionary model to investigate stabilising and equalising mechanisms of coexistence by modelling the evolution of mean trait values affecting competition. Ecological dynamics in their model apply a Lotka-Volterra approach in which intrinsic population growth and the strength of competition both depend on a focal trait (Pastore et al., 2021). Evolution is underpinned by a version of the breeder’s equation in which the trait distribution is normal with a fixed variance. Pastore et al. (2021) demonstrated how trait evolution could affect coexistence and identified eco-evolutionary equilibria at which coexistence is predicted. In another study, Yamamichi et al. (2022) adapted MCT to an eco-evolutionary framework in two competing species. More specifically, they derived expressions for niche overlap and competitive differences assuming single species dynamics go to an equilibrium point and that evolution is rapid enough for traits to stabilise prior to invasion such that both resident and invader are adapted to competition with the resident.

One problem that arises when extending MCT invasibility criteria to integrate evolution is that the ecological dynamics of species densities must be coupled to evolutionary dynamics of traits, which results in a higher-dimensional system. In this case, there are two challenges to determining whether species will coexist: First, determining whether a species can invade when rare in this context is a non-trivial task because it is not obvious a priori at what trait values we should evaluate invasibility. Second, it is not clear whether mutual invasibility is sufficient to imply species coexistence (e.g. in three-species rock-paper-scissors dynamics).

Hence, a more precise analysis of this issue is needed. Here, we do this using the simple two-species competition model analysed by Pastore et al. (2021) and using the framework of persistence theory (Garay, 1989; Schreiber et al., 2011; Patel & Schreiber, 2018), in which all species must persist given initially positive densities. We first characterise the eco-evolutionary equilibrium of the resident species and show that, depending on the parameter values (in particular the trait variances of each species), the invader can evolve to different trait values when rare in this model. We then analyse the potential for coexistence to find precise conditions that ensure coexistence, in the sense of persistence, for any initial trait values. We then highlight that the usual notion of persistence may be too restrictive and not biologically meaningful. To address this, we define a modified form of coexistence, termed *nearly persistent*, which restricts to a subset of initial trait values, and we give precise conditions for this weaker notion of coexistence. Finally, we explore and categorize the possible eco-evolutionary dynamics numerically using the desolve package in R (Soetaert et al., 2010). All code is available on GitHub (Lyberger et al., 2025).

## 2 A two-species eco-evolutionary model

We consider a model of two co-evolving and competing species. Individuals within a species are characterised by an underlying trait *z* that influences their fecundities and competition with conspecifics and heterospecifics. We track changes in population abundance *n*_*i*_(*t*) and trait distribution *ϕ*_*i*_(*z, t*) for species *i*. Note that species have potentially different trait distributions. As in Pastore et al. (2021), we assume that the trait distribution for each species is Gaussian with a dynamic mean 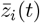 and a fixed variance *V*_*i*_,

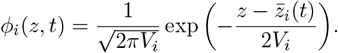

### 2.1 Fecundity and competition

The reproduction and death of an individual are determined by the individual’s intrinsic growth function and its mortality due to intraspecific and interspecific competition. Following Pastore et al. (2021), we assume that the intrinsic growth function of individuals with phenotype *z* in species *i* is *b*_*i*_(*z*), which is defined by a quadratic function,

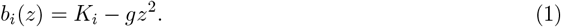

For simplicity, we use the term “fecundity” to denote the function *b*_*i*_(*z*), although this more accurately represents the net growth rate in the absence of competition. The maximum fecundity of an individual in species *i* is *K*_*i*_, which is achieved at the optimal trait value *z* = 0, and *g* determines how quickly fecundity declines when deviating from this optimum. Following Pastore et al. (2021), we can interpret 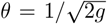 as the width of the fecundity function, with larger values of *g* reflecting a more narrow function and therefore steeper declines when deviating from the optimum (see Figure 1). Equation (1) corresponds to an approximation, for large values of *θ*, of the Gaussian growth function with variance *θ*^2^ assumed in many classical models of character displacement (Roughgarden, 1976; Slatkin, 1980; Taper & Case, 1985; Case & Taper, 2000; Day, 2000), although many of these earlier models assume that the carrying capacity, rather than the fecundity, is a Gaussian function of the trait.

**Figure 1.**
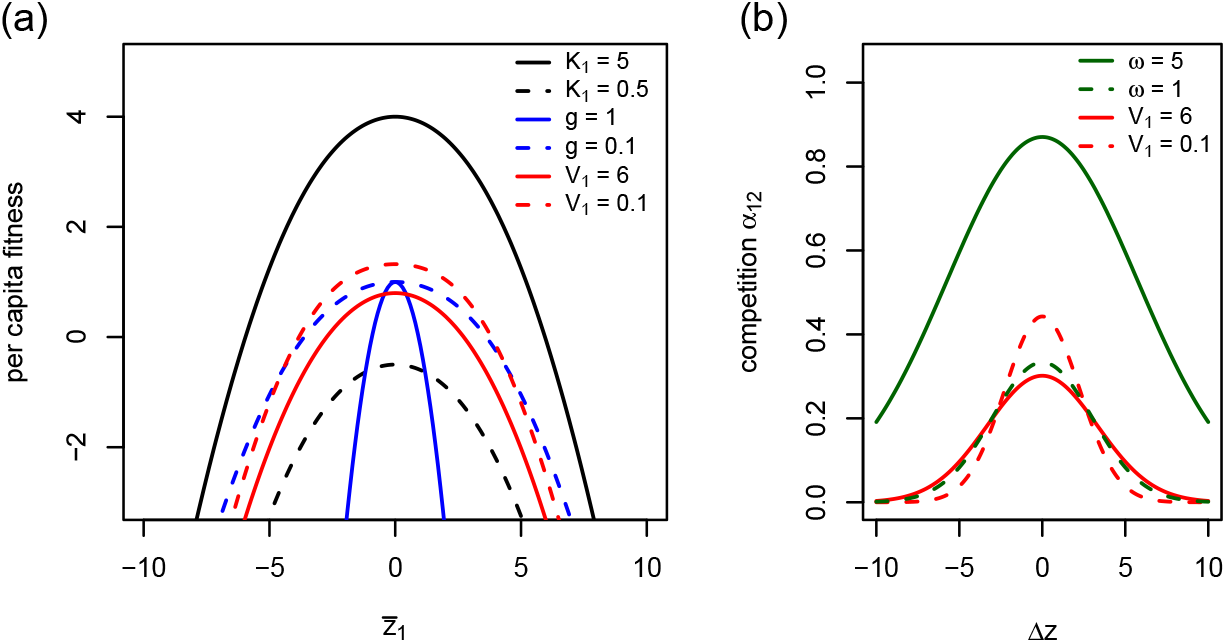
(a) The effect of high values (solid lines) and low values (dashed lines) of *K*_1_, g, and *V*_1_ on invader per capita fitness as a function of the invader’s trait when invader density is 0 and (b) the effect of *V*_1_, and *ω* on competition as a function of the difference between resident and invader traits. Parameters are as follows unless specified in the panel: *ω*=1, g=0.125, *K*_1_ = 2, *K*_2_ = 1, *V*_1_ = *V*_2_=4.

As in Pastore et al. (2021), we assume that competition between two individuals with traits *z*_1_ and *z*_2_ is symmetric and given by a Gaussian kernel,

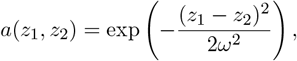

where *ω* is the competition width. Consequently, competition is maximal when individuals have similar trait values (*a*(*z, z*) = 1), and decreases as the absolute difference between traits *z*_1_ and *z*_2_ increases.

### 2.2 Eco-evolutionary dynamics

Using these model ingredients, the joint dynamics of the density and mean trait of species *i* can be written as (Pastore et al., 2021),

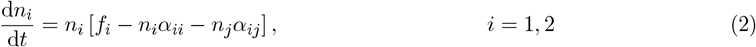

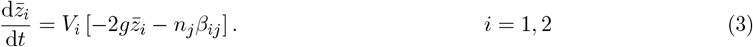

Note the term in brackets of equation (2) is the per-capita growth rate of species *i*, and the term in brackets of equation (3) is the per-capita growth rate gradient with respect to 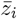. For simplicity, equation (3) implicitly assumes that the trait is fully heritable.

Here, *f*_*i*_ and *α*_*ij*_ are the average fecundity and competition of species *i*, respectively, and can be calculated by integrating the fecundity function and competition kernel over the trait distributions of the species Pastore et al. (2021). Integrating *b*_*i*_(*z*) over the species’ trait distribution *ϕ*_*i*_(*z, t*) defines the average per-capita fecundity of species *i* in the absence of competition,

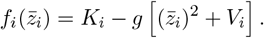

Similarly, the average competition between *i* and *j* is,

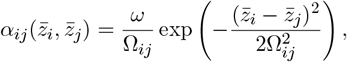

where

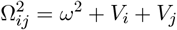

captures the effective competition width, which depends on the intrinsic competition width *ω* and on the trait variances of each species. The average intraspecific competition *α*_*ii*_ is obtained as the case when *i* = *j*, leading to *α*_*ii*_ = *ω/*Ω_*ii*_ where 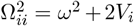 because 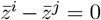 and *V*_*i*_ = *V*_*j*_. The average fecundity (*f*_*i*_) and competition (*α*_*ii*_ and *α*_*ij*_) therefore depend on the trait means and variances of the two species.

Lastly, we calculate the selective effect of inter-specific competition (*β*_*ij*_ in equation (3)) by integrating over the trait distributions *ϕ*_*i*_(*z, t*) and *ϕ*_*j*_(*z, t*), which leads to,

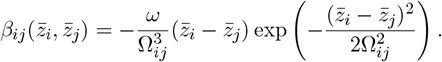

We refer the reader to Pastore et al. (2021) for further details on the derivation of these equations. Apart from a few diffences in notation and the assumption of fully heritable traits, our model is strictly identical to theirs.

### 2.3 Model assumptions

The state space of interest for the model (equations 2-3) is 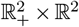. Additionally, our model includes parameters, *K*_1_, *K*_2_, *V*_1_, *V*_2_, *ω*, and *g*, which are all assumed to be positive. Finally, we assume that *K*_*i*_ − *gV*_*i*_ *>* 0 for *i* = 1, 2. If this condition is reversed, then species *i* would not persist even in the absence of competition, which is not a scenario of interest.

Solutions of our model equations are well-behaved in the sense that population densities and trait values are (eventually) bounded from above and thus do not diverge. More precisely, we have the following two propositions:

#### Proposition 1.

*Let* 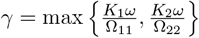. *Then, solutions to (2-3) for any initial condition satisfy*

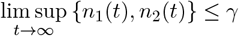

To prove this, first observe that whenever 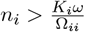, the per-capita growth of species *i* is bounded above by a negative quantity,

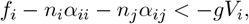

for *i* ≠ *j*. Therefore, all solutions eventually enter 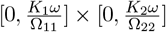.

#### Proposition 2.

*There is a z*^*^ *such that solutions to (2-3) for any initial condition, satisfy*

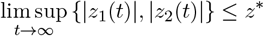

The proof of the second proposition is in the appendix. Altogether, these two propositions give us the desired property of having *dissipative* model equations. That is, there is a compact set 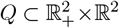 that all solutions eventually enter and remain within. In biological terms, the intraspecific competition within each species and the stabilizing selection around zero ensure that the densities and trait values retain biologically realistic values and do not blow up.

## 3 Resident-invader dynamics

Equations (2) and (3) capture the joint dynamics of densities and mean traits for the two competing species. Pastore et al. (2021) used these equations to calculate how the classical metrics of MCT (niche overlap and the ratio of competitive differences) change over time. However, the key criterion used to derive these metrics in MCT is invasion from rarity. In this section, we first characterise the eco-evolutionary dynamics of the resident (species *j*) in the absence of the invader (species *i*). Then, we characterize the invader trait dynamics according to equation 3. In particular, we show that the invader’s trait may evolve to either a zero or non-zero equilibrium.

### 3.1 Equilibrium of the resident species

#### Proposition 3.

*Solutions to equations (2-3) with n*_*i*_(0) = 0, *n*_*j*_(0) *>* 0 *satisfy*,

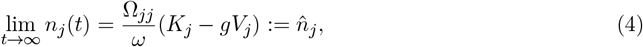

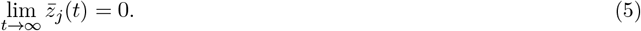

We give a proof of this proposition in the Appendix. This proposition states that the resident species *j* will converge to an equilibrium density 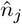 and a mean trait of zero in the absence of a competitor. That is, the mean trait value of species *j* converges to its optimal trait value without interspecific competition 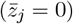. Note that, because 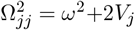, the equilibrium density of species *j* is non-monotonic with respect to the variance of its trait distribution, *V*_*j*_. In the Supplementary Figure 1, we show one example of how the equilibrium population density of the resident depends on its trait variance *V*_*j*_.

### 3.2 Eco-evolutionary invasion dynamics

The fundamental ingredients of MCT are the growth rates of a rare invading species in an environment determined by the resident species. However, when considering eco-evolutionary dynamics, the growth rate of the invading species depends on the distribution of both resident and invader trait values, which are constrained by evolutionary dynamics. In our case, the trait distribution of the invading species *i* is fully described by the mean trait, 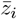, for a fixed value of the trait variance *V*_*i*_. Hence, in the context of eco-evolutionary dynamics, we need to ask: for what mean trait values of the resident and invading species should we test invasion? In the following, we show that the mean trait of the invader approaches an equilibrium point, but that this is not necessarily unique, depending on parameter values.

#### Proposition 4.

*Let* 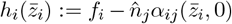 *be the per-capita growth rate of species i at the resident equilibrium of species j given in equations (4-5). Let* 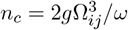. *Then*,

1. *If* 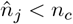, *then*
  a. *h*_*i*_ *is unimodal, with a maximum at* 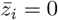.
  b. *solutions to equations (2-3) with n*_*i*_(0) = 0, *n*_*j*_(0) *>* 0 *satisfy* 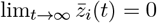
2. *If* 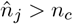, *then*
  a. *h*_*i*_ *is bimodal, with a minimum at* 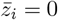 *and maxima at* 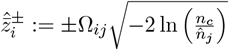.
  b. *solutions to equations (2-3) with n*_*i*_(0) = 0, *n*_*j*_(0) *>* 0 *and* 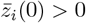 *satisfy* 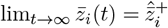

We give a proof of this proposition in the Appendix. This proposition identifies two different outcomes for the invader’s evolutionary dynamics, depending on whether the resident’s equilibrium density is above or below a threshold 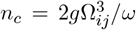. These two outcomes differ by (1) the shape of the invader’s per-capita growth rate at the unique resident equilibrium, in particular the number of peaks (this is characterised by parts 1a and 2a of Proposition 4), and (2) the dynamics of the invader’s mean trait, which can converge either to zero or to non-zero equilibrium values. Biologically, this corresponds to a trade-off between maximising intrinsic growth rate (at 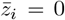) and avoiding interspecific competition, which depends on the population density and trait dynamics of the resident. Hence, the invader’s actual growth rate is not necessarily maximised at a trait value of zero.

Which of these conditions is met depends on the strength of stabilising selection towards the fecundity optimum, *g*, on the competition width, *ω*, on the trait variances in each species (*V*_*i*_ and *V*_*j*_), and on the intrinsic fecundity of the resident *K*_*j*_. When the resident’s population density is higher than *n*_*c*_ (e.g. due to a high intrinsic growth rate, *K*_*j*_, or low trait variance, *V*_*j*_), condition 2 is met and the per-capita growth functions 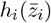 are bimodal (Case A, B and C in Figure 2i). In this case, an invading species with an initial positive mean trait will converge to a fixed positive value 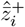, which is distinct from the optimum of the fecundity function at zero (and by the symmetry of our model, if the invading species has an initial negative mean trait, then this will instead converge to 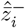). The invader’s per-capita growth rate can be positive at all equilibria (Case A in Figure 2), positive only at 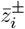 but not at the 0 trait equilibrium (Case B), or negative everywhere (Case C).

**Figure 2.**
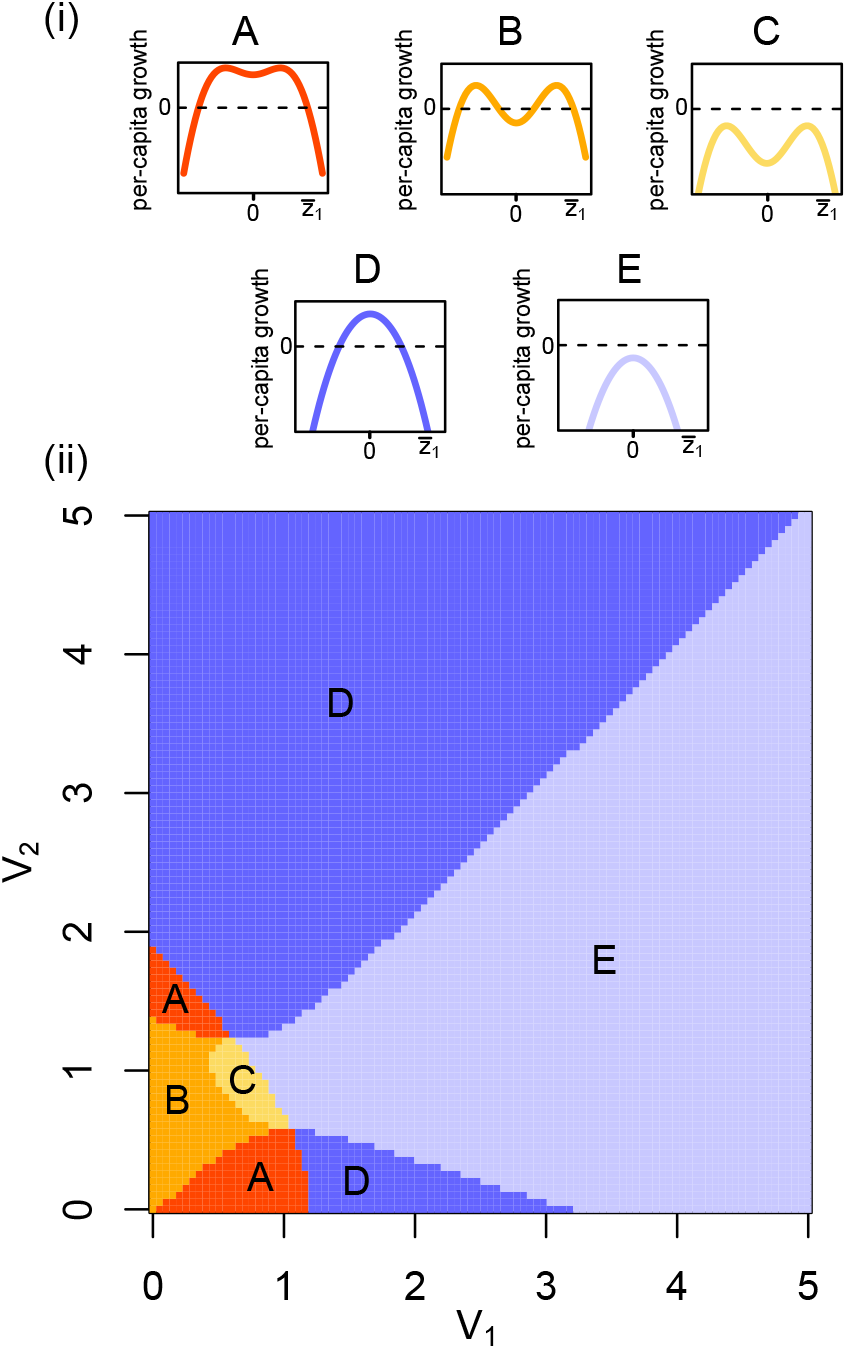
The top panels (i) display the per-capita growth rate function of an invader *i* at equilibrium of the resident, 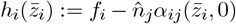. In A, B, and C, condition 2 in proposition 4 is met and the function is bimodal, while in D and E condition 1 is met and the function is unimodal. The bottom panel (ii) shows the effect of variance of the invader *V*_1_ and of the resident *V*_2_ on the outcome of a single species invasion. Parameters are as follows unless specified in the panel: *w* = 1, *g* = 0.125, *K*_1_ = 0.79 and *K*_2_ = 0.80.

On the other hand, when the resident’s equilibrium density is below the threshold *n*_*c*_ (e.g. due to a low intrinsic fecundity, *K*_*j*_, or high trait variance, *V*_*j*_), condition 1 is met and the per-capita growth function 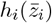 is unimodal (Case D and E in Figure 2i). In this case, the invading species with an initial positive mean trait will converge to a value of zero. The per-capita growth can be either positive everywhere (case D) or negative everywhere (case E). Interestingly, the evolution of invading species *i* does not depend on its maximal fecundity *K*_*i*_.

Figure 2(ii) shows how the variances of the invading and resident species lead to each of these 5 invasion cases, and hence, affects the invasion of the rare species. We observe that the variance of the resident has two opposing effects. On one hand, it may make invasion (positive per-capita growth of the invader) more likely, since it puts a load on the resident, decreasing its equilibrium population density (see equation (4)). On the other hand, it may make invasion less likely since it also decreases intraspecific competition of the resident, thereby potentially increasing its equilibrium population density. In the Supplementary Figure 2, we show an analogous figure to Figure 2(ii) for different values of *K*_1_ and *K*_2_.

To shed light on these results, it is interesting to rewrite the threshold density *n*_*c*_ as

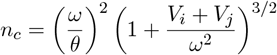

where 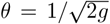 is the width of the fecundity function. Hence, in the absence of trait variances, the threshold *n*_*c*_ corresponds to the square of the ratio of competition width over resource width that characterises whether selection is stabilising or disruptive at the resource optimum in classical character displacement models (Slatkin, 1980; Taper & Case, 1985; Day, 2000; Sasaki & Dieckmann, 2011). Non-zero trait variances increase the threshold *n*_*c*_, but an increase in the invader’s variance *V*_*i*_ only affects the denominator of the condition 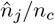 (making it larger), while an increase in the resident’s variance *V*_*j*_ affects both numerator and the denominator. In addition, an increase in the resource width *θ* leads to a lower threhold *n*_*c*_, which means that the condition 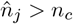 is more easily satisfied. High values of *θ* lead to small values of the ratio *ω/θ*, which corresponds to the condition for the coexistence of more than one type in Gaussian character displacement models (Roughgarden, 1972; Slatkin, 1980; Sasaki & Dieckmann, 2011).

The motivation behind Proposition 4 is similar to the approach proposed by Yamamichi et al. (2022). If we assume that the invader is sufficiently rare that it does not affect the dynamics of the resident population (*n*_*i*_ ≪ *n*_*j*_), but sufficiently abundant that we can use a deterministic gradient equation to describe the change of the mean trait in the invader’s subpopulation, we can assume that the trait value of the invader will be near its equilibrium value. Then, we can examine invasion at these equilibrium values as a proxy for when the invader is rare. However, unlike in Yamamichi et al. (2022), our analysis shows that the equilibrium trait value at which invasion needs to be evaluated is not necessarily unique. Proposition 4 also extends previous results by Case & Taper (2000) who, using a related model, characterised the resident-invader dynamics and showed that the invader’s growth rate could be either unimodal or multimodal depending on a threshold condition (their Equation (17)), which is similar to our condition 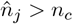. Proposition 4 provides a rigorous demonstration and characterisation of the locations of the peaks, and generalises the results of Case & Taper (2000) to more realistic scenarios where the two species do not have equal trait variances.

## 4 Persistence and Nearly Persistence theorems

In ecological models, coexistence is typically evaluated by the criteria of mutual invasion (e.g., Chesson, 2000b; Barabás et al., 2018; Chesson, 2018; Yamamichi et al., 2022). That is, coexistence is satisfied if each species in the community can invade when its density is low and the rest of the community is at its equilibrium state (e.g., a competitor is at its carrying capacity). While this idea of coexistence is useful, it does not necessarily guarantee that all species are expected to persist and becomes more nuanced in higher-dimensional dynamics. For example, in three competing species, rock-paper-scissor dynamics, in which a single rare species is able to invade a resident equilibrium, it is possible that not all species coexist (see the recent paper by Ranjan et al., 2024 for more description). Persistence theory offers a rigorous approach to establishing coexistence and ensures that population densities of all species eventually stay above some minimal threshold given that they all initially are present (Schreiber, 2006). In this section, we provide precise conditions for the two species to coexist in the sense of persistence given their dynamics in the model written above.

### 4.1 Persistence

We begin with the precise definition of persistence:

#### Definition 1.

*Our model (2-3) is* persistent *if there is a β >* 0 *such that for all initial conditions*

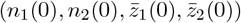

*with n*_1_(0) *>* 0 *and n*_2_(0) *>* 0, *we have that*

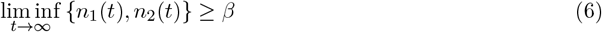

The following result gives sufficient conditions for persistence

#### Theorem 1.

*If for each i* = 1, 2 *and j* = 1, 2 *with j* ≠ *i*

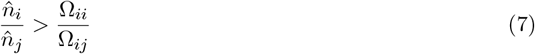

*or equivalently*,

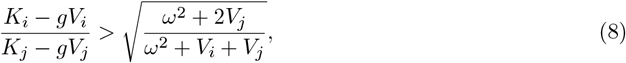

*then our model is persistent*.

We provide a proof of this theorem in the Appendix.

We first observe that when *V*_*i*_ = *V*_*j*_, condition (8) simplifies to *K*_*i*_ *> K*_*j*_ which cannot be simultaneously satisfied by both species. In this case, the species with greater intrinsic fecundity will exclude the other. Differences in variance are therefore a necessary (but not sufficient) condition for persistence.

When *V*_*i*_ ≠ *V*_*j*_, we note that there are really two embedded conditions in condition (8) (for *i* = 1, *j* = 2 and for *i* = 2, *j* = 1). This can be rewritten as,

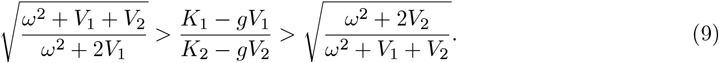

Note that, in this condition, the invasion is evaluated when both 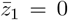 and 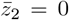. At this value, the resident is at its best trait; it is maximising its intrinsic growth rate. In the case that the invader has a unimodal fitness curve (cf proposition 4), this is where the intrinsic growth of the invading species is also maximised, despite the increased competition it faces with the resident. In the case that the invader has a bimodal fitness curve, this is the worst case scenario for the “invader”; interspecific competition is maximised and the species will not evolve away from this local minimum (i.e., it is “stuck” at this trait value). This makes it impossible to increase invader per capita growth rate at a trait mean that differs from the resident. Hence, if the invader can invade here, then it can eventually invade starting from any initial mean trait, and it is the only condition that we need to check.

It is interesting to ask what happens if the persistence conditions are not met. The following statement shows such a case with a partial converse statement.

#### Theorem 2.

*If*

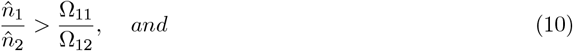

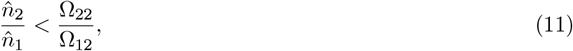

*then the system is not persistent and in particular, solutions to equations (2-3) with* 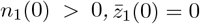 *and* 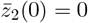 *satisfy* lim_*t*→∞_ *n*_2_(*t*) = 0 *and* 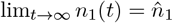.

This captures the case when species 1 is able to invade when rare, but species 2 is not able to invade when rare, when both have an initial mean trait of zero. In this case, we see a dynamic analogous to the classical competitive exclusion of species 2.

The proof is fairly straightforward. It relies on recognizing that the set {(*n*_1_, *n*_2_, 0, 0)|*n*_1_ ∈ ℝ_+_, *n*_2_ ∈ ℝ_+_} is invariant under the model equations. Then, the eco-evolutionary model equations reduce down to the classical Lotka-Volterra ecological equations of two competing species. The conditions given have an equivalence to the classical conditions of species 1 competitively excluding species 2.

### 4.2 Near persistence

In eco-evolutionary dynamics, persistence, classically defined, can be a somewhat restrictive criteria for thinking about coexistence; it requires that each species can invade for *all* initial trait values. In our model, this includes the zero equilibrium mean trait value of the invader, even when it is a local minimum of a bimodal fitness curve. In this case, the trait value of zero is unstable (in the dynamics restricted to the space where *n*_2_ = 0) and small perturbations would allow the invading species to evolve to a different equilibrium value. Hence, it could be the case that the invader is able to invade and subsequently coexist with the resident for some given initial values of its mean trait, i.e. trait-dependent persistence. Because of this, we introduce a less restrictive (but weaker) notion of persistent, called “nearly persistent”, that may be more relevant biologically.

#### Definition 2.

*Our model is* nearly persistent *if there is a β >* 0 *such that for all initial conditions*

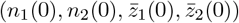

*with* 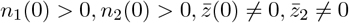, *we have that*,

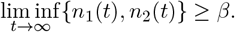

The key distinction is that now we restrict the initial conditions of the trait values to be non-zero.

Hence, we are getting persistence that is dependent on the initial trait values.

The next result provides a sufficient condition for our model to be nearly persistent.

#### Theorem 3.

*If, in addition to (10-11), for each i* = 1, 2 *and j* = 1, 2 *with j* ≠ *i, we have*,

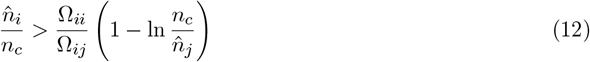

*or equivalently*,

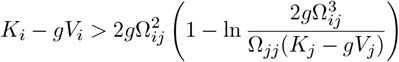

*then our model is nearly persistent*.

We provide a proof of this theorem in the Appendix. Biologically, this statement focuses on the case where each species has a bimodal fitness curve when rare. It only requires invasion of the rare species at the maxima of the bimodal fitness curve when rare, rather than at trait values of 0, i.e., the local minimum of the fitness curve when rare.

Caution is needed to exclude the possibility of eco-evo cycling, in which the population densities of each species get increasingly close to zero even though they eventually invade. This can happen as follows: suppose species 1 is resident, then species 2 invades at its positive trait value and drives species 1 close to zero. When species 1 is rare, species 2 evolves to a trait of zero and species 1 then evolves to a positive trait value, which then leads species 1 to invade. Then, at this trait, species 1 might begin to exclude species 2, and species 2 can get very close to zero density, and the cycle continues.

This theorem states that invasion at fitness peaks is indeed sufficient for near persistence and we do not get these types of cycles.

## 5 Categorization of Dynamics and Numerical Solutions

In this section, we extend beyond persistence to examine the other possible dynamic flows in this model. More specifically, based on pairwise trait-dependent invasion conditions of the invader with a single-resident population, we make predictions on and categorize the two species eco-evolutionary outcomes. These are illustrated in Figure 3. We then test these conditions numerically. We first distinguish three cases based on the modality of the per-capita growth function of the invader species (i.e., when it has density zero). Then, we partition each case by the possible invasion conditions (i.e, whether growth is positive or negative at the equilibra), which leads to a total of 15 possibilities. The three cases and conditions for the per-capita growth function modality are as follows:

**Figure 3.**
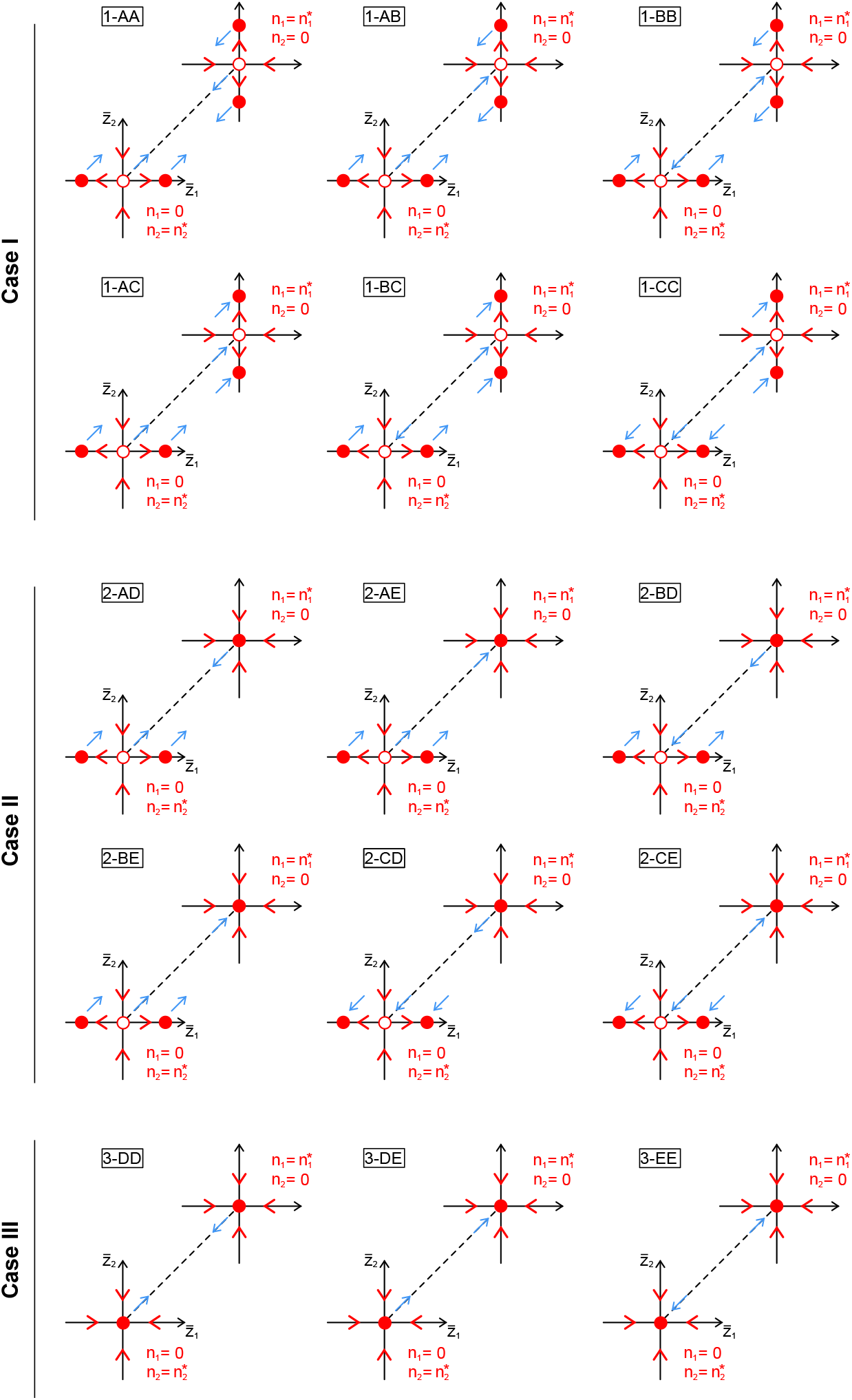
Two-dimensional representation of four-dimensional space in which the system’s dynamics operate. In the front: visualizing the evolutionary dynamics of species 1 is at zero and species 2 is at equilibrium. In the back: visualizing the evolutionary dynamics when species 2 is at zero and species 1 is at equilibrium. Stable and unstable equilibria points (with respect to the trait dynamics on the single-species resident invariant set) are depicted with filled and unfilled circle symbols, respectively. The dotted line indicates the invariant set formed when 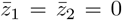. See section 5 for a detailed description of Case I, II and III. Each of these three cases can then be combined with the cases A, B and C described in Figure 2.

- Case I: If 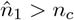 and 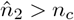. In this case, both species have bimodal per-capita growth functions as invaders when the opposing species is a resident at equilibrium. That is, the trait dynamics of the invader has three equilibra.
- Case II: If 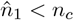 and 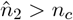. In this case, species 1 has a bimodal per-capita growth function (i.e., three equilibra for its trait dynamics) when species 2 is resident at equilibrium but species 2 has a unimodal per-capita growth function (i.e., unique eq. point at 0), when species 1 is resident at equilibrium.
- Case III: If 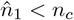 and 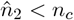. In this case, both species have unimodal per-capita growth functions as invaders when the opposing species is a resident at equilibrium. That is, the trait dynamics of the invader has a unique equilibra at 0.

We further partition these cases based on the flow near the boundary. When an invading species has bimodal per capita growth rate curves, we categorize the boundary flow into three relevant categories (labeled A, B, C in Figure 2), while when an invading species has unimodal growth curves, we categorize into two relevant categories (labeled D and E in Figure 2). In (A) and (D), the invading species can invade at all trait equilibria points. In (B), the invading species can only invade at the per capita growth peaks. In (C) and (E), the invading species cannot invade at any trait equilibria.

Altogether, the 3 cases and these flows near each boundary result in the 15 possibilities shown in Figure 3. There are 6 possibilities within Case I, since each species can exhibit one of three invasion categories (A, B, or C). Similarly, there are 6 possibilities within Case II and 3 possibilities within Case III. From numerically examining the 15 possibilities, we observe that there are five qualitatively distinct ecological scenarios possible in this model, including persistence:

1. Persistent: the two competing species will coexist for any initial mean trait values provided both are initially present in the community.
2. Nearly persistent: the two competing species will coexist for when either species has non-zero initial mean trait values, provided both are initially present in the community. However, if both species start with an initial trait of zero, then one species will be excluded.
3. Competitive exclusion: species *i* will competitively exclude the species *j* for any initial trait values. Species *j* will not evolve to coexist.
4. Near competitive exclusion: species *i* will competitively exclude species *j* if either species has a non-zero initial mean trait value. If both species have an initial mean trait value of zero, then there will be a priority effect.
5. Priority effect: for any initial mean trait values of each species, one species will exclude the other, depending on their initial densities.

Which ecological scenario results in which case of the boundary dynamics and flow is summarized in Table 2. We find that the precise eco-evolutionary outcome we predict strongly depends on the variances and maximal fecundity of the two species. Figure 4 demonstrates how assuming equal species variances results in near persistence at small variances and priority effects at large variances, whereas a large asymmetry in variances leads to competitive exclusion, where the species with the smaller variance wins. Finally, a small asymmetry in species variances often results in persistence. When maximal fecundity values *K* differ between species, eco-evolutionary outcomes tend to vary, and we do not observe regions of priority effects. In the Supplementary Figure 3, we show analogous figures to Figure 4 for different different values for *K*_1_ and *K*_2_.

**Table 1:**
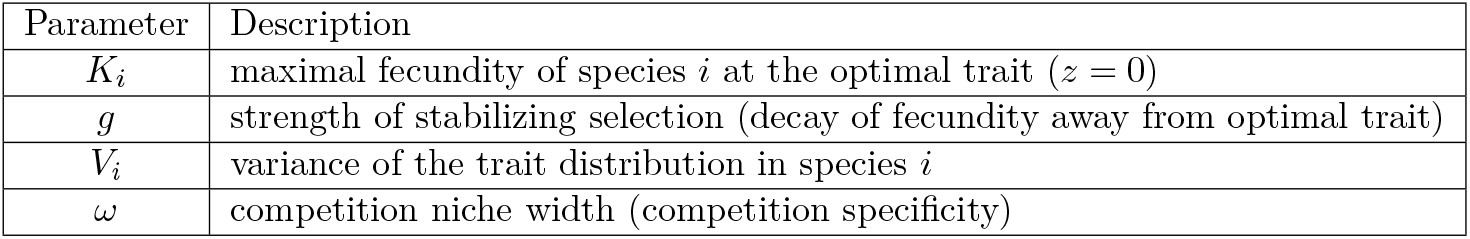
Description of key parameter values used in a model of eco-evolutionary dynamics between two competing species.

**Table 2:** Predictions of qualitative behaviors from boundary dynamics and flow near boundary. Letters A-E refer to the per-capita growth rate functions of species 1 and 2 as invading species as shown in Figure 2. Number in parentheses (i.e., I-III) indicates the boundary dynamic cases (described in the main text). Legend: P = Persistent, NP = Nearly Persistent, CE = Competitive Exclusion, NCE= Near Competitive Exclusion, PE = Priority Effect.

|  |  | Species 1 |  |  |  |  |
| --- | --- | --- | --- | --- | --- | --- |
|  |  | A | B | C | D | E |
| Species 2 | A | (I) P | (I) P | (I) CE | (II) P | (II) CE |
|  | B |  | (I) NP | (I) NCE | (II) NP | (II) NCE |
|  | C |  |  | (I) PE | (II) CE | (II) PE |
|  | D |  |  |  | (III) P | (III) CE |
|  | E |  |  |  |  | (III) PE |

**Figure 4.**
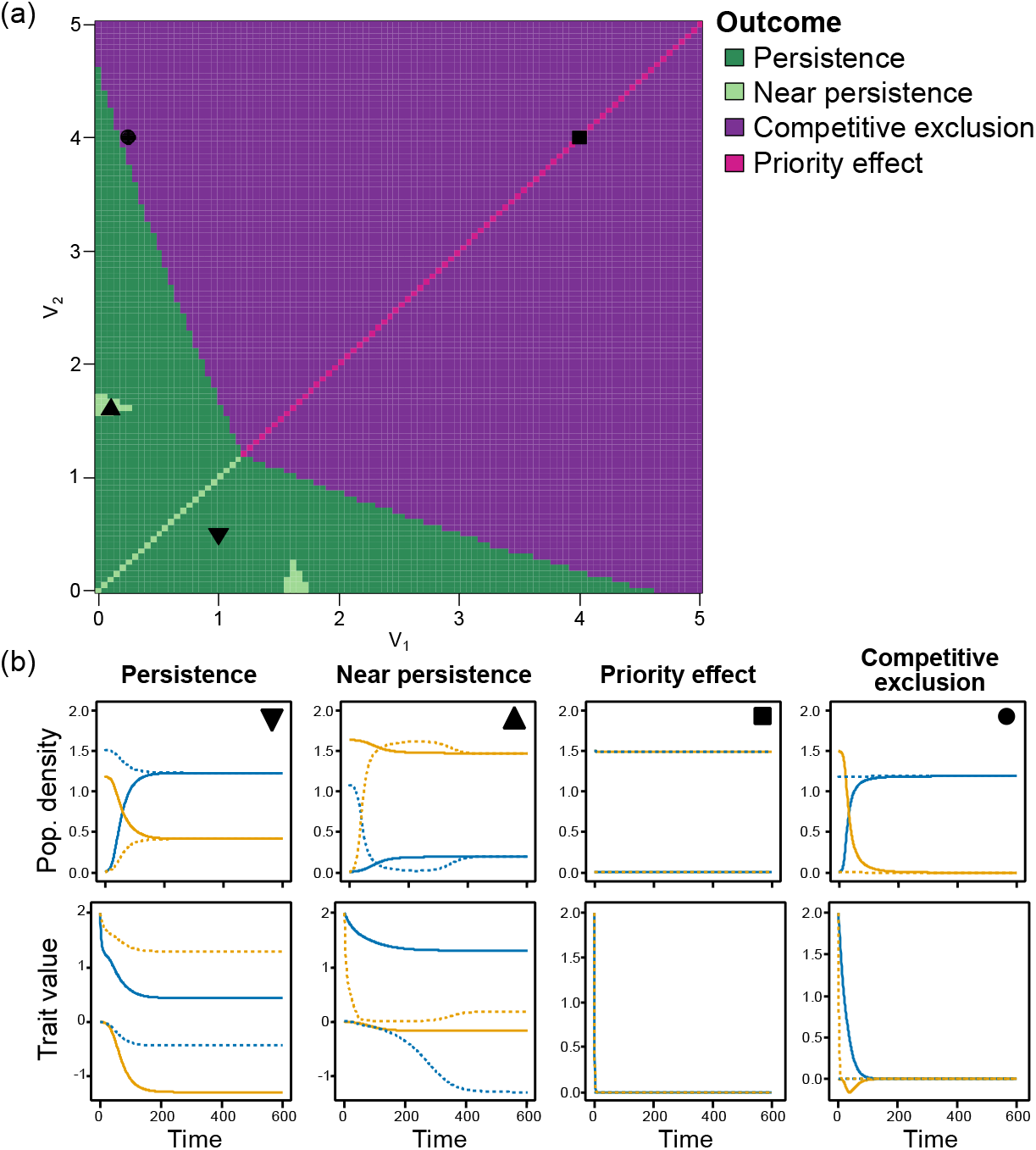
(a) The effect of changing variance in both species on the qualitative behaviour produced. Parameters are as follows: *w*=1, *g* = 0.125, *K*_1_ = *K*_2_ = 1. The black point represents the values from Pastore et al. 2021. (b) Eco-evolutionary temporal dynamics simulated for three invasion scenarios. Each column shows two stacked panels: the upper panel depicts population densities *n*_*k*_(*t*) and the lower panel depicts mean trait values 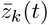 for two competing species. Solid lines represent the invasion of species 1 (blue) into a resident population of species 2 (yellow), whereas dashed lines reverse the invasion order. Parameters common to all simulations are competition width *w* = 1, intrinsic growth–rate width *g* = 0.125, and carrying capacities *K*_1_ = *K*_2_ = 1. The scenarios differ only in the standard deviations of the Gaussian competition kernels (*V*_1_, *V*_2_): *case 1AA, persistence* (1, 0.25); *case 2DB, nearly persistence* (0.1, 1.6); *case 3DD, priority effect* (4, 4); *case 3DE, competitive exclusion* (0.25, 4). Simulations were run for 600 time units with numerical output every 1 unit. Symbols in panel (b) illustrate simulations with corresponding parameter values in panel (a).

## 6 Discussion

Modern coexistence theory (MCT) has been a powerful framework for understanding coexistence of species in communities (Chesson, 2000b; Barabás et al., 2018; Chesson, 2018), spatial environments (e.g., Chesson, 2000a; Ellner et al., 2022), and fluctuating environments (e.g., Chesson, 2017; Ellner et al., 2019; Johnson & Hastings, 2023). A critical assumption of MCT, however, is that species traits are fixed and trait variance is ignored. Here, we have expanded on previous attempts to incorporate evolutionary dynamics into MCT (Pastore et al., 2021; Yamamichi et al., 2022) to understand how trait evolution and intraspecific trait variation affect species coexistence. We find that (1) with ecoevolutionary dynamics, the choice of species traits is not arbitrary when evaluating mutual invasion conditions for an invader and resident species, but determined by the dynamics of the species’ mean traits, (2) depending on the trait variances in each species, there can be multiple trait values at which the invasion condition needs to be evaluated to accurately assess the possibility of coexistence, and (3) limitations of the classic invasion criteria of MCT can be overcome by evaluating conditions for persistence. This leads to a more nuanced picture of the role of invasion analyses in determining coexistence.

Our first result is that the evolutionary dynamics of the resident and invader species (e.g. the change in each species’ mean traits) determine the trait values at which the classical MCT invasion conditions need to be evaluated. In our 2-species model with quadratic resource function and Gaussian competition kernel, the equilibrium values of the mean traits for the resident and invader can be analytically calculated for a fixed combination of the trait variances. We show that, when the resident density is below a threshold, the invader’s trait converges to the optimum trait value for the resident, so that the outcome of the invasion is solely determined by the difference in trait variances in the two species. On the other hand, when the resident density is above a threshold, the invader’s trait can converge to either a positive or negative equilibrium value. Overall, this allows us to identify five qualitatively different patterns for invasion, depending on the shape of the per-capita growth rate of invader, which can be either a unimodal or bimodal function of the trait. Our results therefore show that incorporating evolutionary dynamics into MCT is not a straightforward task. Previous studies (e.g., Yamamichi et al., 2022) have emphasised the need to estimate the invasion potential of species at a specific eco-evolutionary (EE) point determined by the coupled eco-evolutionary dynamics. However, the possibility that there could be multiple EE points was not recognised, although earlier character displacement models had hinted at this possibility (Case & Taper, 2000).

Classical MCT relies on invasion analyses by calculating the mean growth rate of an invader against the backdrop of a resident species or community at equilibrium (Barabás et al., 2018). Coexistence is achieved if every species in the community has a positive growth rate when its density is rare. This framework has been particularly successful and widely applied to 2-species communities using the niche overlap and ratio of competitive differences metrics (Barabás et al., 2018; Yamamichi et al., 2022). Persistence theory (Schreiber et al., 2011) provides an alternative approach to analysing coexistence, which is more robust in higher-dimensional communities. The second contribution of our study is to apply persistence theory to our eco-evolutionary model to better characterise coexistence of the two species in the sense of persistence. Persistence has a rich history developed in the late 1970s (Freedman & Waltman, 1977; Gard & Hallam, 1979), and early 1980s (Gard, 1980, 1981, 1982; Freedman & Waltman, 1984), but has been limited in the context of eco-evolutionary dynamics (but see Schreiber & Patel, 2015; Patel & Schreiber, 2018; Browne & Yahia, 2023). Hence, applying persistence theory to eco-evolutionary models in the classical notion of persistence requires that population densities eventually remain bounded from below by a positive density *for all* initial trait values. However, in some cases, this might not be the most biologically relevant criterion. Here, we present a weaker notion of persistence: restricting persistence to a subset of initial starting trait values, we can show that the population densities of all species are eventually bounded from below by a positive density.

On the technical side, the proof of our near persistent theorem uses previous results from Garay (1989). Roughly speaking, Garay (1989) considered ecological models on a state space in which the boundary of the space is invariant for that model. Typically, the boundary is the extinction set (in which at least one species in a community has density zero). They provided conditions for showing the this set is repelling flow under the model. We recognized that their proof could be applied to other invariant sets and, provided this *includes* the extinction set, then we can still show this weaker notion of persistence. Further investigations of persistence in models that include an evolutionary component and how persistence may depend on traits would be worthwhile to build more understanding of how evolution impacts coexistence.

Beyond these technical advancements, our analysis emphasises the key role of the trait variances of each species in determining coexistence. In contrast to earlier studies that relied on a similar competition model (e.g., Case & Taper, 2000; Pastore et al., 2021), we allow for the two species to have distinct trait variances and show that relaxing the assumption that trait variance is the same in both species critically affects the conditions for invasion and persistence. Given identical trait variances, the model of Pastore et al. (2021) predicts either near persistence (given small variances) or priority effects (given large variances; Figure 3a). Instead, our model predicts either persistence or competitive exclusion in the more realistic case where species trait variances differ. In addition, we show that 15 distinct cases of qualitative phase portraits can be distinguished based on the initial conditions and trait variance of species. This emphasises that a rigorous extension of MCT to incorporate evolutionary dynamics requires a careful consideration of the dynamics of the eco-evo system, even in a simple twospecies competition model.

In empirical systems, interacting species will naturally differ in their intraspecific trait variances, with profound implications for the rate of evolution of each species and for the possibility of species coexistence. In an eco-evolutionary model focused on effects of intraspecific variation, Barabás & D’Andrea (2016) found that intraspecific trait variances generally limit the scope for coexistence, except when species have similar mean trait values but differ highly in trait variances. Further investigation of the role of trait variances is therefore needed to elucidate how asymmetry in trait means and variances affects the coexistence of interacting species.

Intraspecific trait variation is central to rapid trait change, and both can affect ecological processes and interactions (e.g., Thompson, 1998; Bolnick et al., 2011; Lankau, 2011; Kopp & Matuszewski, 2014; Cope et al., 2022). The invasion analysis we develop, as the one proposed by Yamamichi et al. (2022), relies on an implicit assumption that the density of the invader is much smaller than the trait variance, so that trait evolution occurs on a fast time scale while the invader is still rare. This is the opposite of the classical assumption of MCT, which assumes trait variance to be so small that evolution is negligible and species traits are fixed. A careful analysis of what happens in between these two extreme scenarios is still needed. In addition, species trait variances are unlikely to remain constant as selection can also be expected to alter the shape of the trait distributions. If species trait variances could instead evolve, intraspecific variation would likely decrease as a consequence of interspecific competition (Barabás et al., 2022a). This might make persistence more likely in our model because we generally find persistence at lower species trait variances (Figure 4). Evolution of trait variances might therefore expand the range of initial conditions that result in persistence.

We hope that this analysis supports further investigations into how the persistence of species may depend on their traits. In particular, in this model, we assumed the fecundity was symmetric around the optimal trait value (quadratic function). Asymmetric fitness functions could lead to even more nuances in trait-dependent persistence. That is, invading species may be able to invade at some trait values but not others, and hence, coexistence may be more critically dependent on the underlying evolutionary dynamics. Additionally, our model assumes trait distributions remain Gaussian even under the impacts of selection. Although this is a very classical assumption in quantitative genetics, it is natural to expect that selection impacts the full trait distribution, leading to non-Gaussian distributions (e.g. skewed or multimodal distributions). One way to better capture the dynamics of the full distribution would be to examine the dynamics of the variance under selection, while assuming higher order moments remain constant. Finally, here we consider arguably the simplest case of two competing species. Many species of interest live in more complex communities involving several species competing as well as other types of interactions. In these cases, it is natural to predict that there may be even more potential evolutionary outcomes dependent on initial traits. Hence, thinking about the notion of invasion when rare in an eco-evolutionary context may require even more care.

## Acknowledgements

This manuscript was supported by joint funding between the French Foundation for Research on Biodiversity (FRB) Centre for the Synthesis and Analysis of Biodiversity (CESAB) and the German Centre for Integrative Biodiversity Research (sDiv). It was written as part of the Unification of Modern Coexistence Theory and Price Equation (UNICOP) project. LG was supported by the German Science Foundation (DFG, project No. 511084840). KL was supported by the NSF Postdoctoral Research Fellowships in Biology Program (No. 2208947). SP was supported by the Oregon State University College of Science Research and Innovation Seed (SciRIS) grant. VL was supported by the Conselleria d’Innovació, Universitats, Ciència i Societat Digital – Generalitat Valenciana (Project: CIGE/2023/16).

## Appendix A: Proofs

We will use standard techniques from dynamical systems and, more specifically, persistence theory, to show the results given in the main text. The proofs rely heavily on the main results from Garay, 1989 and Patel & Schreiber, 2018 on persistence in ecological systems. These require dissipative systems and so we begin with the proof that our model is dissipative.

### A.1 Proof of Proposition 2

We will show that the mean trait value for species 1 is eventually bounded, under our model equations. The same proof applies for species 2. The equation for the trait dynamics of a species 1 is

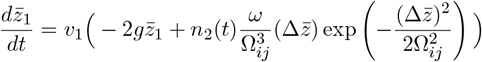

where 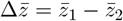

- First, observe that 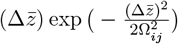 is bounded. That is, there is a *δ* so that for all 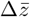, we have

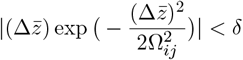
- By proposition 1 eventually in time *n*_2_ is bounded by *γ*. Hence, there is a *T* such that for all *t > T*, we have 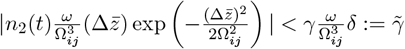
- Finally, define 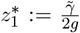. Then, there is a *β >* 0 such that for all *t > T* and 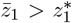, we have 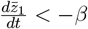 (and if 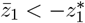, then 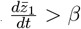). This implies there is a 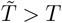 such that for all 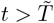, 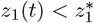.

We can apply the same procedure to yield a 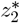. Then let 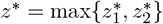.

### A.2 Terminology

For the remaining proofs, we will first set up some necessary terminology. As in Patel & Schreiber, 2018, our eco-evolutionary model equations can be expressed through the general ecological system with feedback equations:

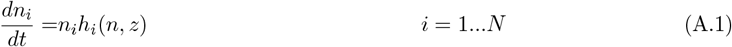

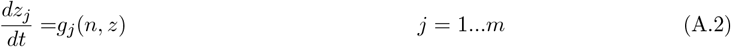

where *n*_*i*_ is the population density of species *i* and *z*_*j*_ is a feedback variable.

Let 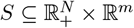 be the state space and 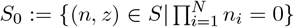 be the extinction set of this space. Assuming existence and uniqueness of solutions, let Φ : ℝ*× S* → *S* be the flow defined by A.1-A.2 and 𝒪 (*y*) = {Φ(*t, y*)|*t* ≥ 0} is the forward orbit for initial condition *y* = (*n, z*) ∈ *S*. Assuming dissipativity, there is a compact set *Q* ⊂ *S* so the forward orbits for any initial condition *y* ∈ *S* eventually enter and stay in *Q*.

The *ω*-limit set of *y* ∈ *S* is 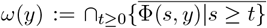 and the *α*-limit set of *y* ∈ *S* is 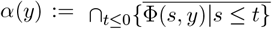. [Note that here *ω*-limit is used for identifying a set, which is distinct from the parameter *ω*.] The *ω*-limit set of a set *Y* ⊂ *S* is 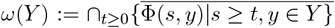. A set *Y* ⊂ *S* is invariant if {Φ(*t, y*)|*t* ∈ ℝ, *y* ∈ *Y*} = *Y*. The global attractor is the maximal compact invariant set *ω*(*Q*).

For an invariant set Γ, we say that *z* ∈ Γ is chain accessible from *y* ∈ Γ if for every *ϵ >* 0, *t >* 0, there is an (*ϵ, t*)-chain from *z* to *y*, i.e., a sequence {*y*_1_ = *y, y*_2_, …, *y*_*n*+1_ = *z*; *t*_1_, *t*_2_, …, *t*_*n*_} with *y*_*i*_ ∈ Γ, *t*_*i*_ *> t* and *d*(Φ(*t*_*i*_, *y*_*i*_), *y*_*i*+1_) *< ϵ* for *i* = 1, …, *n*. A point *y* ∈ Γ is called chain recurrent if it is chain accessible to itself. We let *R*_*C*_(Γ) be the chain recurrent set. Finally, the set Γ is chain recurrent if every *y* ∈ Γ is chain recurrent.

A collection of sets ℳ = {*M*_1_, *M*_2_, …, *M*_*ℓ*_} is a Morse decomposition for a compact invariant set Γ if the sets are pairwise disjoint, isolated, invariant, compact sets, so that for every 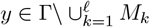 there are integers *i*(*y*) and *j*(*y*) with *i < j* and *ω*(*y*) ⊂ *M*_*i*_ and *α*(*y*) ⊂ *M*_*j*_.

### A.3 Proof of Proposition 3-4

Without loss of generality, we will set *i* = 1 (invader) and *j* = 2 (resident). Under the hypothesis *n*_1_ = 0, the model equations become

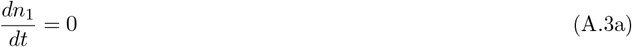

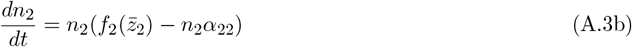

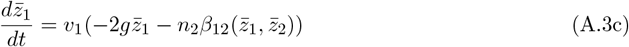

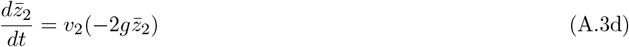

The proof relies on recognizing that these are hierarchically decoupled: the flow of 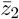 only depends on 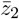, the flow of *n*_2_ depends on 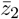 and *n*_2_ and the flow of 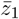 depends on *n*_2_ and 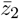. We will provide proofs for each claim made in the propositions

- Equation (A.3d) is a simple linear differential equation with exponential decay solutions 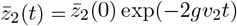. Hence, 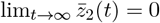.
- Using the solution of equation (A.3d), observe that equation (A.3b) becomes an asymptotically autonomous differential equation

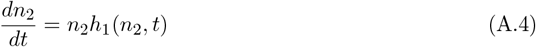

where

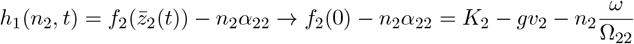

where the convergence is uniform in ℝ. We can make statements about the asymptotically autonomous flow in relation the the autonomous flow, which in this case is

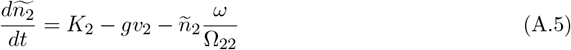

The chain recurrent set of (A.5) is the set 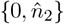, where 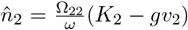. Then, Theorem 1.8 of Mischaikow et al (1995) gives us that the *ω*-limit of *n*_2_(0) for (A.4) is contained in 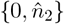. Finally, if *n*_2_(0) *>* 0 then there is an *ϵ*_1_ *>* 0 and *T*_1_ *>* 0 so that *n*_2_(*t*) *> ϵ*_1_ for all *t > T*_1_. To see this, observe that by the assumption *K*_2_ *> gv*_2_, we have that there is a *T*_2_ *>* 0 and *ϵ*_2_ *>* 0 so that *h*_1_ *> ϵ*_2_ for all *n*_2_ *< ϵ*_1_ and *t > T*_2_. Hence, if *n*_2_(0) *>* 0 then 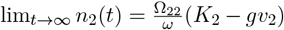.

- In a similar fashion, equation (A.3c) is

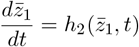

where

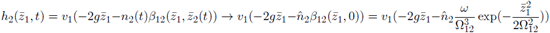

and again this convergence is uniform in ℝ. We define the autonomous differential equation

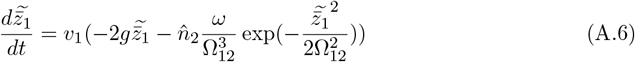

The flow defined by (A.6) has two possibilities for its chain recurrent set:

1. if 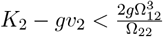, then the chain recurrent set is {0}

2. if 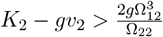, then the chain recurrent set is 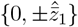

By a similar argument to above, we can show that in case 2 and 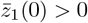, the non-autonomous flow defined by (A.3c) has 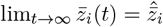

### A.4 Proof of Theorem 1

The statement is:

If for each *i* = 1, 2 and *j* = 1, 2 with *j* ≠ *i*

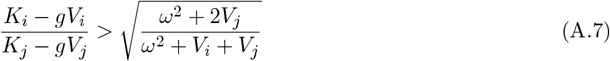

then our model is permanent.

We will use the following theorem (from Patel & Schreiber, 2018) to prove this statement.

#### Theorem 4.

*Let* ℳ = {*M*_1_, *M*_2_, …, *M*_*ℓ*_} *be a Morse decomposition for S*_0_ ∩ Γ *where* Γ *is the global attractor for A.1. If, for each M*_*k*_ ∈ ℳ, *there exists a p*_*k*1_, *p*_*k*2_, …, *p*_*kn*_ *>* 0 *such that for every y* ∈ *M*_*k*_, *there is a T*_*y*_ *such that*

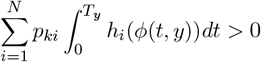

*then A.1 is persistent*.

This theorem extends persistence theory results of purely ecological systems to account for feed-backs such as the evolutionary one we consider in this paper. In our equation,

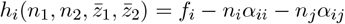

The proof proceeds in two parts. First, we identify an appropriate Morse decomposition for our system. The Morse decomposition depends on which case we consider. Then, we will show, for each Morse set, the existence of a 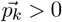 to satisfy the inequality in Theorem 4.

- For Case I, we define the Morse decomposition as the following set of ordered equilibrium points: 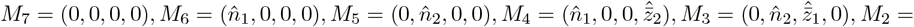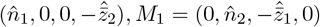. (These are partially illustrated in Figure 3.)
- Next, observe that the inequality in Theorem 4 for two species is

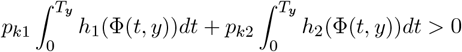

For each set *M*_*k*_ and *z* ∈ *M*_*k*_, we let 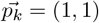 and *T*_*y*_ = 1. We show this inequality is met. (In fact, any positive vector and any *T*_*y*_ *>* 0 would do.) Then, recognizing that each Morse set is an isolated equilibrium point, the condition, for Morse set *M*_*k*_ becomes

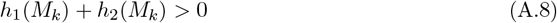

- For *M*_7_, the standing assumptions that *K*_*i*_ − *gv*_*i*_ *>* 0 for *i* = 1, 2 ensure inequality A.8 is met.
- Furthermore, for evenly indexed Morse set, *M*_*k*_, we have *h*_1_ = 0 and hence, need only check

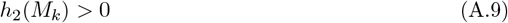

For oddly indexed Morse sets, except *M*_7_, we have *h*_2_ = 0 and need only check

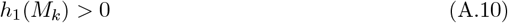

Hence, this persistent condition reduces to checking invasion (positive growth) of an invader *j* at Morse sets with purely resident species *i*.

- For *M*_6_ and *M*_5_, the hypothesis of the theorem ensures that *h*_2_(*M*_6_) *>* 0 and *h*_1_(*M*_5_) *>* 0.
- Next, observe that if *h*_2_(*M*_6_) *>* 0, then *h*_2_(*M*_4_) *>* 0 and *h*_2_(*M*_2_) *>* 0. This is true since *α*_12_(0, *z*) ≤ *α*_12_(0, 0) for all *z* ∈ ℝ.
- An analogous argument holds to show that if *h*_1_(*M*_5_) *>* 0, then *h*_1_(*M*_3_) *>* 0 and *h*_1_(*M*_1_) *>* 0.
- Finally, the proof for these Cases II and Case III follows similarly to Case I.

### A.5 Proof of Theorem 3

To show the result on near persistence, we use an earlier persistence result from Garay, 1989. Garay used the Ura-Kimura Theorem, which roughly states that if the stable set of a compact isolated invariant set is contained in itself, then it is asymptotically unstable. Here, the stable set is

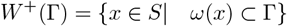

Garay used this theorem to give persistence conditions by showing the boundary of a locally compact metric space is unstable.

#### Theorem 5

(Garay). *Let* (*E, d*) *be a locally compact metric space and let S be a closed subset of E and ϕ a dissipative dynamical system for which the boundary ∂S is invariant (and* Γ *is a maximal compact invariant set). Further, assume that for each component C of the chain recurrent set R*_*C*_(Γ ∩ *∂S*), *we have*

1. *a γ >* 0 *such that* {*x* ∈ *S\∂S*|*d*(*x, C*) *< γ*} *contains no entire trajectories*
2. *W* ^+^(*C*) ∩ *S\∂S* = *∅*

*Then, the system is persistent*.

In the proof Garay uses the Ura-Kimura theorem to show the boundary *∂S* is asymptotically unstable, by examining a corresponding chain recurrent set, and that this then implies persistence. Going through the proof, we recognize that the result of asymptotic instability applies more generally to any non-open invariant set, not just boundary sets. Hence, we can apply this proof to show the invariant set 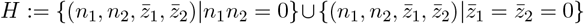 is asymptotically unstable. Here *H* is the union of the extinction set, i.e., where at least one of the species is extinct and the set where both species have mean trait of zero.

To apply this theorem, we focus on the chain recurrent set *R*_*C*_(Γ ∩ *H*). These are the equilibrium points:

- *C*_1_ := (0, 0, 0, 0)
- 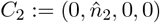
- 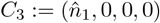
- 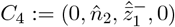
- 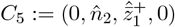
- 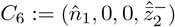
- 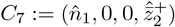

Notice that for each pair of components, *C*_*i*_ and *C*_*j*_, with *i* ≠ *j, C*_*i*_ and *C*_*j*_ are not chain accessible from each other. Next, we show that *C*_1_ is isolated and that *W* ^+^(*C*_1_) ∩ *S\H* = *∅*. Suppose that this is not true. Then, for any *ϵ*−neighborhood around *C*_1_, denoted *B*_*ϵ*_(*C*_1_), there is a *y* ∈ *B*_*ϵ*_(*C*_1_)\ *H* so that *ϕ*(*y, t*) ∈ *B*_*ϵ*_(*C*_1_) for all *t* ≥ 0. From our standing assumptions, there is a *δ >* 0 and *ϵ >* 0 so that *h*_1_(*y*) *> δ* and *h*_2_(*y*) *> δ* for all *y* ∈ *B*_*ϵ*_(*C*_1_) (*h*_1_ and *h*_2_ are the per-capita growth rates as defined in previous proof). Since *y* ∈ *B*_*ϵ*_(*C*_1_)\ *H* implies either that *n*_1_ *>* 0 or *n*_2_ *>* 0, then there is a *t >* 0 such that *ϕ*(*y, t*) ∉ *B*_*ϵ*_(*C*_1_), which is a contradiction. A similar proof holds for *C*_2_, …, *C*_7_. Hence, conditions 1 and 2 of the theorem are met and *H* is asymptotically unstable.

