## Supplementary Figures for "Persistence and near persistence via trait evolution: pathways to coexistence"

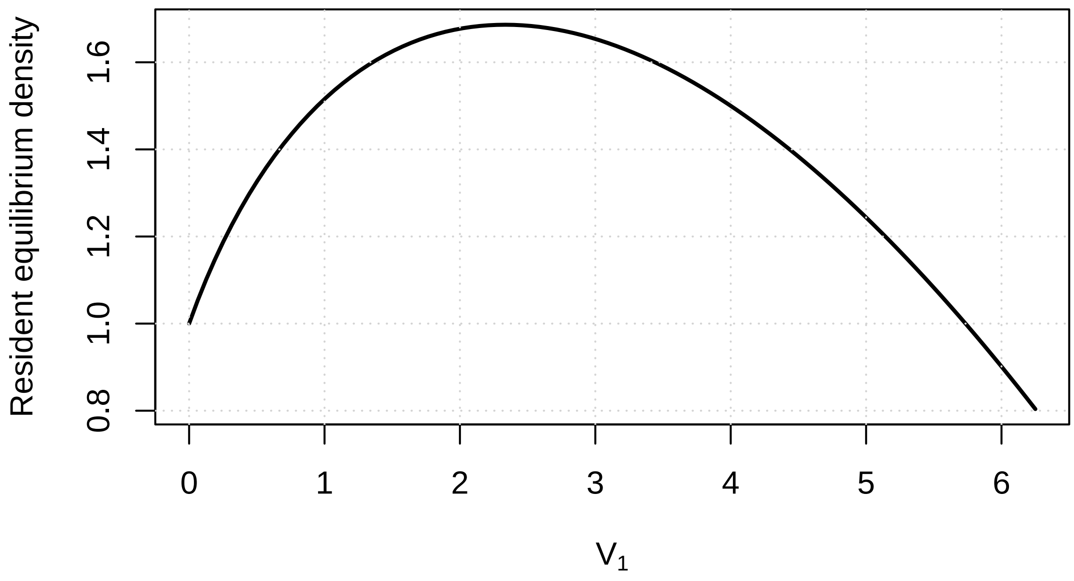


FIGURE S1. Resident equilibrium density of species 1 as a function of its trait variance. The equilibrium density decreases with increasing variance due to increasing stabilizing selection and intraspecific competition. Parameters are as follows: $w = 1$, $g = 0.125$, $K_1 = 1$.

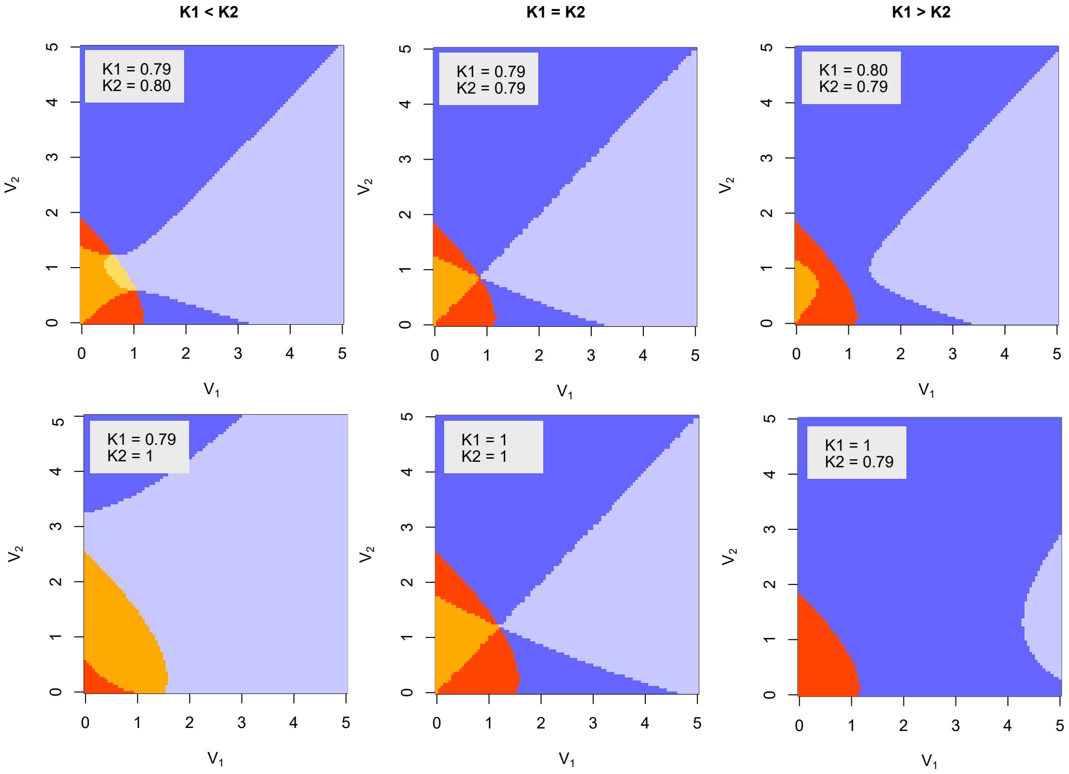


FIGURE S2. Different panels display the effect of variance of the invader $V_1$ and of the resident $V_2$ on the outcome of a single species invasion depending on values of $K$. Parameters are as follows unless specified in the panel: $w = 1$, $g = 0.125$. Colours reference the same cases as in Figure 2i of the main text.

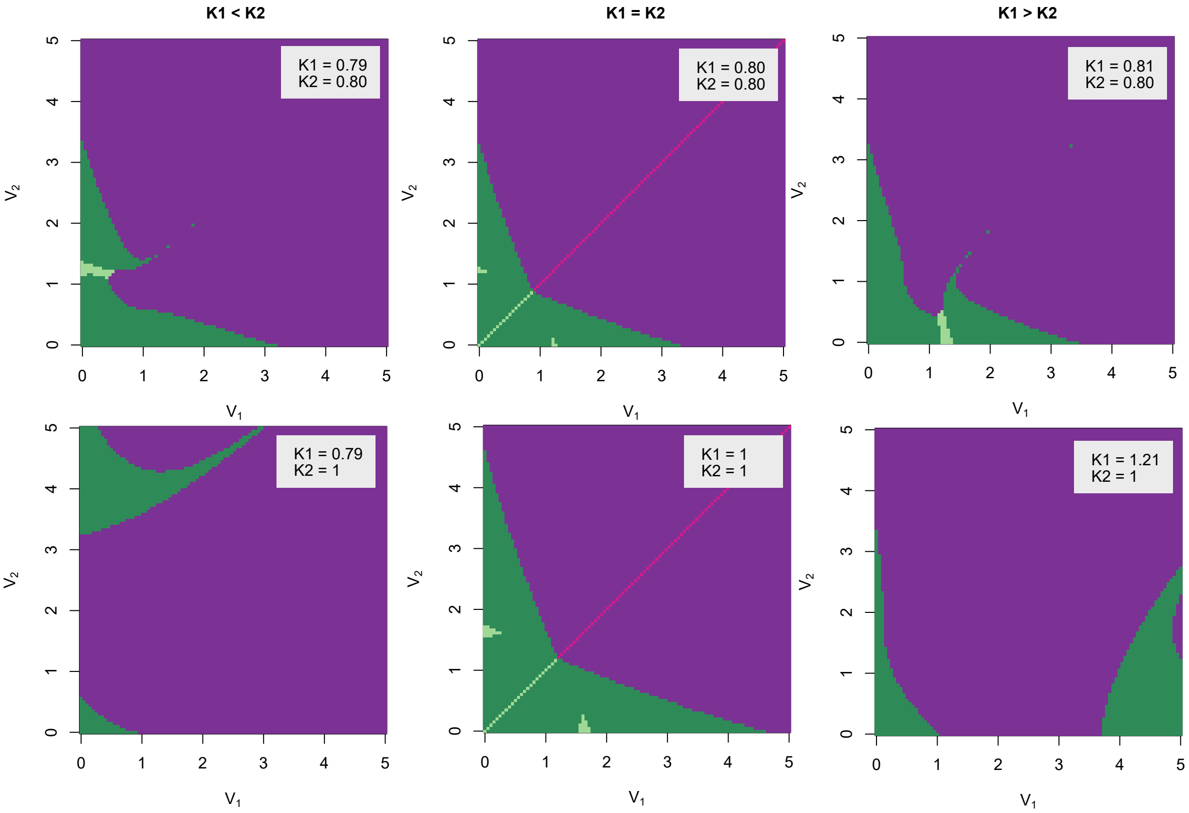


FIGURE S3. The effect of changing variance in both species on the qualitative behaviour produced with varying maximal fecundity values for invader and resident species. Parameters are as follows: $w$=1, $g$ = 0.125 or given in figure panel. The black point represents the values from Pastore et al. 2021. The colors represent the same outcomes as in Figure 5a of the main text.
